# D-alanylation of Teichoic Acids in *Bacilli* impedes the immune sensing of peptidoglycan in *Drosophila*

**DOI:** 10.1101/631523

**Authors:** Zaynoun Attieh, Mireille Kallassy Awad, Agnès Rejasse, Pascal Courtin, Ivo Gomperts Boneca, Marie-Pierre Chapot-Chartier, Vincent Sanchis Borja, Laure El Chamy

**Affiliations:** Unité de Recherche Environnement, Génomique et Protéomique, Laboratoire de Biodiversité et Génomique Fonctionnelle, Faculté des sciences, Université Saint-Joseph de Beyrouth- Liban; Micalis Institute, INRA, AgroParisTech, Université Paris-Saclay, 78350 Jouy-en-Josas, France; Institut Pasteur, Unité biologie et génétique de la paroi bactérienne, 75724 Paris cedex 15, France; INSERM, Equipe Avenir, 75015 Paris, France

## Abstract

Modification of cell wall components is a prominent mean for pathogens to hinder host immune defenses. Here, using the *Drosophila* model, we aimed at characterizing the role of D-alanine esterification of teichoic acids (TAs) in the resistance of *Bacillus thuringiensis* to host defenses *in vivo*. We show that, by preventing cationic antimicrobial effectors-mediated bacterial lysis, this cell wall modification also limits the release of peptidoglycan immunostimulatory fragments thus impeding their sensing and the subsequent induction of the IMD- NF-κB pathway. Interestingly, we show that this strategy is also adopted by *Lactobacillus plantarum*, a *bona fide* commensal, to fine-tune its immunomodulatory potential in the *Drosophila* gut. Markedly, we show that the D-alanylation of TAs is essential for the resistance of *L. plantarum* to gut lysozyme. Altogether our data shed light on the mechanism underlying the persistence and the low immunostimulatory potential of *L. plantarum* in the *Drosophila* gut.

## Introduction

The external surface of microorganisms is a central platform for interactions with multicellular hosts. Evolutionarily selected Pattern Recognition Receptors (PRRs) of the innate immune system sense key microbial surface molecules such as fungal glucans, Gram-negative bacterial lipopolysaccharide (LPS), Gram-positive bacterial Peptidoglycan (PGN) and Teichoic Acids (TAs) to alert the host of the microbial non-self and to initiate immune responses (Medzhitov, 2007). PRRs, such as the Toll-like receptors and NOD-like receptors act at the forefront of the mucosal immune barrier. They control the expression of several immune induced genes such as antimicrobial peptides (AMPs) and cytokines, the differential profiles of which dictate the outcome of the ensued immune response (Hold, Mukhopadhya, & Monie, 2011). Therefore, several hypotheses consider microbial surface molecules as plausible candidates for effector probiotic molecules (Kleerebezem et al., 2010). Likewise, several pathogens have evolved specialized mechanisms to modify their external surface in order to avoid detection or to develop resistance to potent, highly conserved, immune effectors such as AMPs and lysozymes (Anaya-Lopez, Lopez-Meza, & Ochoa-Zarzosa, 2013; Bechinger & Gorr, 2017; Koprivnjak & Peschel, 2011; Ragland & Criss, 2017). In humans, both lysozyme and AMPs are expressed at surface epithelial cells as well as in phagocytes and thus act at the frontline of the host defenses (Dommett, Zilbauer, George, & Bajaj-Elliott, 2005). Most AMPs are small cationic molecules with a pronounced amphipathy, a property that accounts, as commonly agreed, for their membrane targeting microbicidal activity. According to the model, AMPs are preferentially attracted to negatively charged surfaces of the microbial cell envelopes where they get embedded into the hydrophobic regions of the lipid membranes, thereby leading to membrane destabilization and ultimately to cell death (Joo, Fu, & Otto, 2016). Lysozyme comprises an evolutionarily conserved family of bactericidal proteins that are found in species ranging from bacteriophages to man (Dommett et al., 2005). Their canonical killing mechanism relies on their muramidase activity, which cleaves the β-(1,4)-glycosidic bond between N-acetylmuramic acid and N-acetylglucosamine moieties of the bacterial cell wall peptidoglycan. Resulting loss of cell wall integrity leads to rapid cell lysis and death (Callewaert & Michiels, 2010; Ragland & Criss, 2017). The cationic nature of lysozyme is very important for bactericidal activity since it favors its binding to the negatively charged bacterial cell envelop and hence access to the PGN substrate.

In addition to the thick layer of PGN, the cell wall of Gram-positive bacteria is characterized by the presence of highly charged anionic polymers of repeating alditol phosphate residues called Teichoic acids (TAs). TAs represent the most abundant component of Gram-positive bacterial cell wall and play crucial roles in their pathogenesis (Brown, Santa Maria, & Walker, 2013; Joo et al., 2016). Notably, modification of TAs by addition of positively charged D-alanine esters residues reduces their net negative charge thus lowering the attraction of cationic antimicrobial peptides to the bacterial cell wall. This process is performed by the gene products of the *dlt* operon that is highly conserved in most Gram-positive bacteria including probiotic and pathogenic species (Abachin et al., 2002; Abi Khattar et al., 2009; Heaton & Neuhaus, 1992; Kristian et al., 2005; Perego et al., 1995; Peschel et al., 1999). According to currently available genome sequences, the *dlt* operon comprises four to five genes, *dltXABCD* (Abi Khattar et al., 2009; Kamar et al., 2017; Perego et al., 1995). DltA is a D-alanine-D-alanyl carrier protein ligase that catalyzes the D-alanylation of the D-alanyl carrier protein DltC (Debabov et al., 1996; Heaton & Neuhaus, 1992, 1994). The roles of DltX, DltB and DltD are less clear (Debabov, Kiriukhin, & Neuhaus, 2000; Kamar et al., 2017; Neuhaus & Baddiley, 2003). Nevertheless, in all tested Gram-positive bacteria species, genetic disruption of any of the five genes of the *dlt* operon completely abrogates its function. Compared to wild-type strains, *dlt* mutants have a higher negative charge on the cell surface and are significantly susceptible to cationic antimicrobial effectors including AMPs and lysozyme (Fabretti et al., 2006; Kovacs et al., 2006; Kristian et al., 2005; Kristian et al., 2003; Perea Velez et al., 2007). In addition, *in vivo* studies clearly show a significantly attenuated virulence of the *dlt* mutants compared to wild-type strains of many human pathogens (Abachin et al., 2002; Abi Khattar et al., 2009; Fisher et al., 2006; Perego et al., 1995; Peschel et al., 1999; Poyart et al., 2003). However, despite deep interest in the characterization of the function of the *dlt* operon, our understanding of its exact role in the bacterial resistance to the innate immune system *in vivo* remains limited.

*Drosophila* has long emerged as a well-suited model for the study of host-pathogen interactions (Ferrandon, 2013; Lemaitre & Hoffmann, 2007). The hallmark of the *Drosophila* host defense is the immune-induced expression of genes encoding potent AMPs (Ferrandon, Imler, Hetru, & Hoffmann, 2007; Lemaitre & Hoffmann, 2007). These peptides are synthesized by the fat body cells (the fly immune organ) and secreted into the hemolymph where they constitute the main effectors of the humoral systemic response. AMPs are also produced locally at the epithelial barriers such as the gut, where they act synergistically with ROS to limit the infections (Ferrandon, 2013; Imler & Bulet, 2005). Two highly conserved NF-κB signaling cascades, the Toll and the Immune Deficiency (IMD) pathways, control the expression of AMP genes. Whereas both pathways contribute to the regulation of the systemic immune response, only the latter regulates the expression of AMP genes in the gut through the activation of the NF-κB transcription factor Relish (El Chamy, Matt, Ntwasa, & Reichhart, 2015; Ferrandon, 2013). The IMD pathway is elicited upon the sensing of DAP-type PGN fragments by members of the Peptidoglycan Recognition Protein (PGRP) receptors family; the membrane bound PGRP-LC and the intracellular PGRP-LE (Chang, Chelliah, Borek, Mengin-Lecreulx, & Deisenhofer, 2006; Chang et al., 2005; Choe, Lee, & Anderson, 2005; Choe, Werner, Stoven, Hultmark, & Anderson, 2002; Gottar et al., 2002; Kaneko et al., 2004; Kaneko et al., 2006; Leulier et al., 2003; Mellroth & Steiner, 2006; Ramet, Manfruelli, Pearson, Mathey-Prevot, & Ezekowitz, 2002). Potential immune effectors, such as lysozymes are also expressed at the epithelial barrier (Daffre, Kylsten, Samakovlis, & Hultmark, 1994). However, their global contribution to the host defense and intestinal homeostasis remains obscure. More recently, *Drosophila* has become a powerful model for the characterization of the molecular mechanisms underlying the mutualistic host-microbiota interactions (Erkosar & Leulier, 2014; W. J. Lee & Brey, 2013; Ma, Storelli, Mitchell, & Leulier, 2015). Indeed, several studies have proven the relative simplicity of the fly intestinal microbiota, which comprises approximately 30 phylotypes with a major representation of *Lactobacillaceae* and *Acetobacteraceae* (Blum, Fischer, Miles, & Handelsman, 2013; Chandler, Lang, Bhatnagar, Eisen, & Kopp, 2011; Corby-Harris et al., 2007; Cox & Gilmore, 2007; Staubach, Baines, Kunzel, Bik, & Petrov, 2013; C. N. Wong, Ng, & Douglas, 2011). Reports have begun to illustrate the impact of this microbial community on the *Drosophila* host biology (Buchon, Broderick, Chakrabarti, & Lemaitre, 2009; Iatsenko, Boquete, & Lemaitre, 2018; Jones et al., 2013; Sharon et al., 2010; Shin et al., 2011; Storelli et al., 2011; A. C. Wong, Dobson, & Douglas, 2014). Notably, the gut commensals largely influence the *Drosophila* midgut transcriptome and promote the expression of genes associated with gut physiology and metabolism as well as tissue homeostasis and immune defenses (Broderick, Buchon, & Lemaitre, 2014; Erkosar & Leulier, 2014). Interestingly, some *Lactobacillus plantarum* strains recapitulate the benefits of a *bona fide Drosophila* commensal thus providing a simple model for the investigation of the intricate mechanisms underlying the impact of intestinal bacteria on host physiology and mucosal immunity (Jones et al., 2013; Storelli et al., 2011). In this context, high throughput analyses have shown that the gut-associated bacteria set an immune barrier in the *Drosophila* gut through the activation of a transcriptional program that is largely dependent on the IMD pathway (Bosco-Drayon et al., 2012; Broderick et al., 2014; Buchon et al., 2009; Ryu et al., 2008). Remarkably though, the expression of AMP genes remains at its basal level compared to that induced upon an oral infection (Bosco-Drayon et al., 2012) and any alteration in this transcriptional profile leads to commensal dysbiosis, which dramatically affects the host fitness and lifespan (Bonnay et al., 2013; Broderick et al., 2014; Ryu et al., 2008). Recent studies have shown that the immune response is compartmentalized in the *Drosophil*a gut (Bosco-Drayon et al., 2012; Buchon, Broderick, & Lemaitre, 2013) and that multiple negative regulators intervene at different levels of the IMD pathway in order to fine-tune its activation (Aggarwal & Silverman, 2008; Bischoff et al., 2006; Guo, Li, & Lin, 2014; Kleino et al., 2008; Lhocine et al., 2008; Mellroth, Karlsson, & Steiner, 2003; Paredes, Welchman, Poidevin, & Lemaitre, 2011; Ryu et al., 2004; Zaidman-Remy et al., 2006). Among these negative regulators are catalytic members of the PGRP family, such as PGRP-SC and PGRP-LB, which degrade PGN fragments into non-immunostimulatory moieties and the intracellular protein, Pirk, which binds to the PGRP-LC and PGRP-LE receptors thus disrupting downstream signaling complex (Aggarwal et al., 2008; Kleino et al., 2008; Kleino et al., 2017; Lhocine et al., 2008). However, similarly to infectious bacteria, *L. plantarum* and the major elements of the fly microbiota produce DAP-type PGN and most of the IMD negative regulators act in a feedback loop. Thus, what makes this microbial community only a mild inducer of the AMP genes and how it is tolerated in the *Drosophila* gut remains intriguing (Bosco-Drayon et al., 2012).

Here, using the *Drosophila* model, we aimed at a better characterization of the intricate *dlt*-dependent mechanisms that allow bacteria to resist the innate immune response *in vivo*. Our results show that, D-alanylation of TAs impedes the sensing of PGN from *B. thuringiensis* in *Drosophila* thus hampering the activation of the IMD pathway upon a systemic infection. This mechanism is also used by *L. plantarum* to fine-tune its induced immune response in the *Drosophila* gut. In particular, we show that the D-alanylation of TAs is essential for the resistance of *L. plantarum* to intestinal lysozyme and thus for its persistence a core component of the *Drosophila* microbiota but also for the modulation of its ensued NF-κB-dependent epithelial immune response.

## Results

### Beyond resistance to AMPs, D-alanylation of TAs in *Bacilli* prohibits the systemic activation of the *Drosophila* IMD pathway

We have recently shown that *B. thuringiensis* is highly virulent to adult *Drosophila* in a septic injury infection model. Using a *dltX* mutant of *B. thuringiensis 407* Cry^−^ strain (*Bt407ΔdltX*), we showed that this phenotype is completely dependent on the D-alanylation of its TAs, which confers resistance to the IMD-dependent humoral immune response (Kamar et al., 2017). Here, we assess whether this phenotype is due to the resistance of *Bt407* to cationic AMPs *in vivo*. For that, we resorted to a rescue experiment of the susceptibility of adult flies to the infections by the wt and the *dlt* mutants of *Bt407 via* the overexpression of *Cecropin*, an-IMD responsive gene encoding a cationic AMP. *Cecropin* overexpression was driven by the UAS-Gal4 system in *Drosophila* using the fat-body specific driver c564-Gal4 (Harrison, Binari, Nahreini, Gilman, & Perrimon, 1995) (Figure 1 – figure supplement 1). In addition to our previously published *Bt407ΔdltX* mutant, our survival experiments employed a full *dlt*-operon deletion mutant, *Bt407Δdlt*, which we generated by allelic exchange with a kanamycin resistance cassette (see experimental procedures). Loss of function of the *dlt* operon was confirmed by quantification of D-Alanine esterified to TAs by HPLC analysis, which proved impaired D-alanylation of TAs in the *Bt407Δdlt* mutant as compared to the wt strain (Figure 1 – figure supplement 2). The survival curves of infected adult flies presented in Figure 1 clearly show that *Bt407* is highly virulent to both wt and *relish* mutant flies whereas the *Bt407ΔdltX* and *Bt407Δdlt* mutants are only pathogenic to the latter (Figure 1 A and C). In addition, the similar susceptibility of wt flies and *relish* mutants to the *Bt407* infection is mirrored by the rate of bacterial growth in the hemolymph of the infected flies (Figure 1 B and D). The bacterial loads are equivalent to those retrieved from *relish* mutants upon an infection with *Bt407Δdlt* and *Bt407ΔdltX* (Figure 1 B and D). These results confirm that the D-alanylation of TAs confers *Bt407* resistance to the systemic humoral immune response in *Drosophila*. Moreover, whereas the overexpression of *Cecropin* does not alter the virulent phenotype of *Bt407*, it significantly ameliorates the survival of the wt flies and completely rescues the phenotype of the *relish* mutants when infected with the *Bt407ΔdltX* and *Bt407Δdlt* (Figure 1A and C). These results correlate with the cessation of bacterial growth in the hemolymph of surviving infected flies (Figure 1 B and D). Altogether, these results indicate that D-alanylation of TAs in *Bt407* confers resistance to Cecropin *in vivo*. This phenotype is strictly dependent on the activity of the *dlt* operon as complementation of the *Bt407ΔdltX* mutant with the *dltX*-ORF (*Bt407ΔdltΩdltX)* completely restores a virulence similar to that of the wt *Bt407* strain (Figure 1 – figure supplement 3).

**Figure 1:**
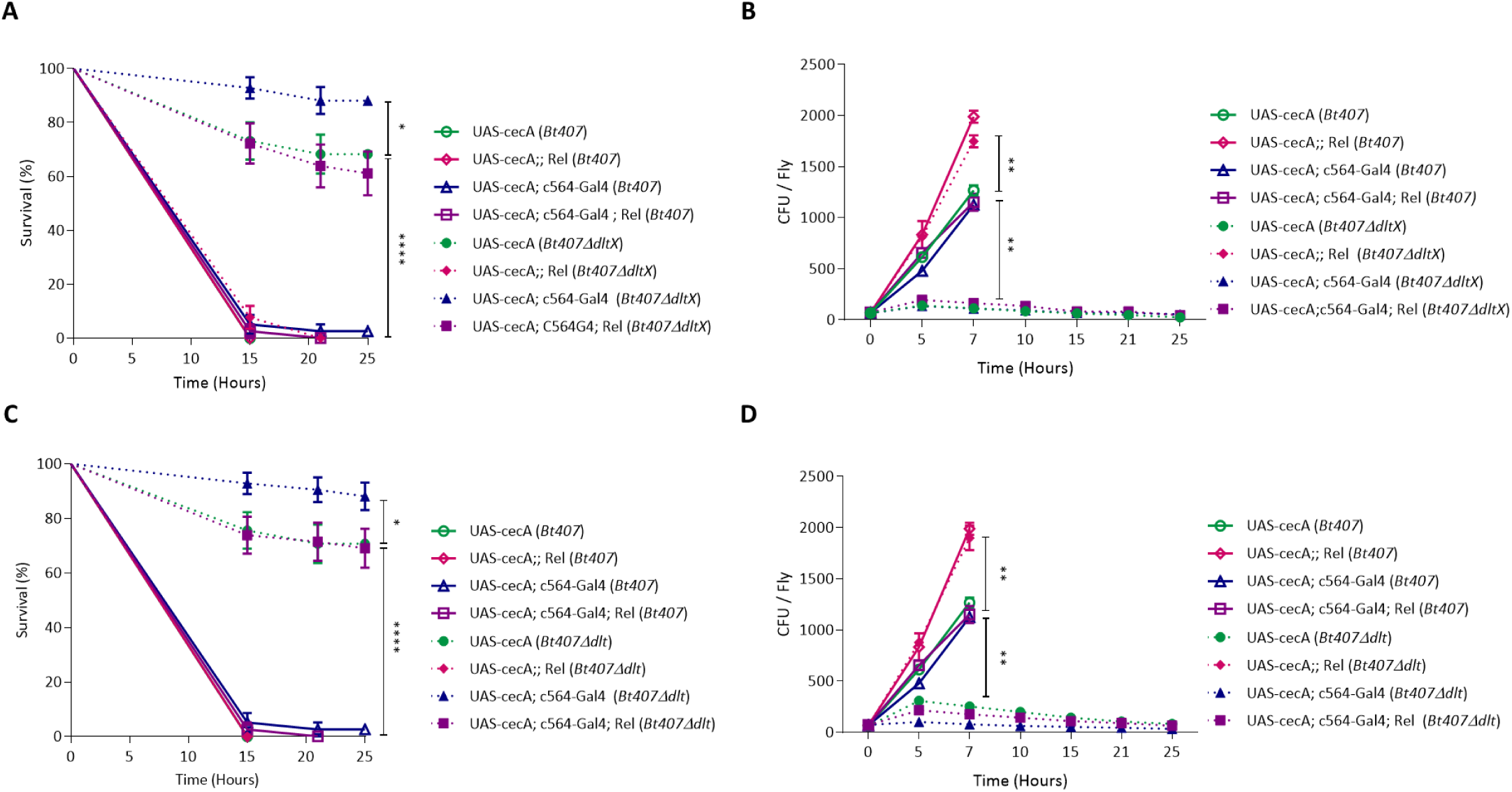
D- alanylation of TAs confers *Bacillus thuringiensis* resistance to the AMP-dependent systemic immune response in *Drosophila*. (A and C) Survival of adult wild-type (wt) or relish (*Rel*) mutant, overexpressing or not a *Cecropin A* transgene (under the control of the c564-Gal4 fat body driver) to an infection with *Bt407, Bt407ΔdltX* (A) and *Bt407Δdlt* (C). (B and D) Internal bacterial loads retrieved from adult flies infected with *BT407*and *Bt407ΔdltX* (B) or Bt407*Δdlt* (D). The Colony Forming Unit (CFU) counting was performed only on survival flies. Data are pooled from three independent experiments (mean and s.d.). Statistical tests were performed using the Log Rank test for the survival assays and Mann-Whitney test for the CFU counting within Prism software (ns: *p* > 0.05; * 0.01< *p* < 0.05; **:0.001< *p* < 0.01; ***: 0.0001< *p* < 0.001; ****: *p* < 0.0001). The exact *p* values are listed in Figure 1 – source data 1.

Since the D- alanylation of TAs conveys a modification of the bacterial surface, we assessed whether it affects the sensing of *Bt* by the *Drosophila* innate immune system. Hence, in order to compare the activation of the IMD pathway by *Bt407* and the Δ*dlt* mutants, we infected the flies by septic injury and quantified the expression of *Diptericin*, an AMP-encoding gene, four hours following infection. The *Bt*- triggered *Diptericin* expression was compared to that induced by *Escherichia coli*, a conventionally used microbial inducer of the IMD pathway (Dushay, Asling, & Hultmark, 1996; Hedengren et al., 1999; Kaneko et al., 2004; Leulier et al., 2003). As shown in Figure 2A, *Bt407* is only a mild-inducer of the IMD pathway as compared to *E. coli*. However, both *Bt407ΔdltX* and *Bt407Δdlt* infection result in *Diptericin* expression level similar to that induced by *E. coli*. This response is strictly dependent on the activation of the IMD pathway since it is abolished in *PGRP-LC* and *relish* mutant flies. The phenotype is completely reverted in the *Bt407ΔdltX* complemented strain (*Bt407ΔdltXΩdltX*) (Figure 2A). Similar results were obtained from another read-out of the IMD pathway, *Attacin D* (Figure 2B). Altogether, our results indicate that beyond resistance to AMPs, the D-alanylation of TAs impedes the sensing of *Bt407* by the innate immune system, thus significantly reducing its immune-stimulatory potential and further contributing to its pathogenicity in *Drosophila*.

**Figure 2:**
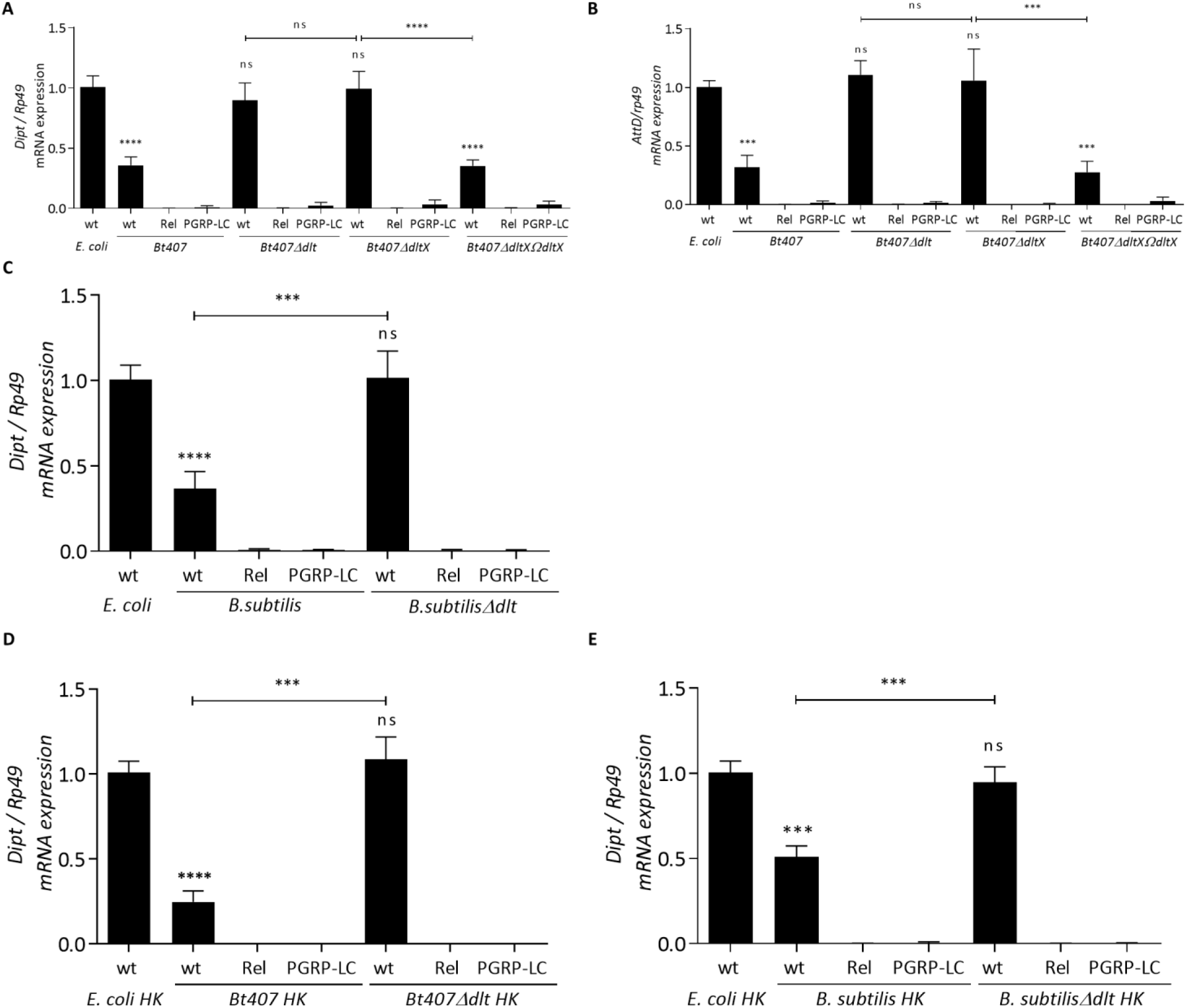
The D- alanylation of TAs confines the induction of the IMD pathway upon a systemic infection of *Drosophila* by *Bacilli*. Relative expression of the *Diptericin* (*Dipt*) (A, C D and E) or *Attacin D* (*AttD*) (B) transcripts in wild-type (wt) or *relish* (*Rel*) mutant flies induced by living or heat-killed *Bt407, Bt407Δdlt, Bt407ΔdltX* and *Bt407ΔdltXΩdltX (A, B and D) or B. subtilis (C and E)*. Transcripts expression was measured by RT-qPCR in total RNA extracts 4 hours upon the induction. Ribosomal protein 49 (Rp49) transcript was used as reference gene. Transcripts levels are compared to that triggered in wt flies infected by *E. coli* as a control. Data obtained from three independent experiments are combined in single value (mean ± sd). Statistical tests were performed using the Mann-Whitney test within Prism software (ns: *p* > 0.05; * 0.01< *p* < 0.05; **:0.001< *p* < 0.01; ***: 0.0001< *p* < 0.001; ****: *p* < 0.0001). The exact *p* values are listed in Figure 2 – source data 1.

Next, we asked whether this mechanism of *dlt*-dependent camouflage of *Bt* is also used by other *Bacilli* species. Therefore, we checked for the immune induction of the IMD response in flies infected with a wt *Bacillus subtilis* (*B. subtilis*) and its *dlt* mutant (*B. subtilisΔdlt*). As shown in figure 2C, the *Diptericin* expression profile elicited by these bacterial strains is equivalent to that activated upon the infection of the flies by *Bt407* and the *Bt407Δdlt* mutant respectively. These data suggest that the *dlt* function results in a restricted elicitation of the IMD pathway also in *Bacilli*. To exclude that the limited induction of the IMD pathway results from its active inhibition from live *Bacilli*, we repeated these experiments using heat killed bacteria and found similar results (Figure 2D and 2E). We conclude that the D-alanylation of TAs highly masks PGN from sensing by PGRP-LC.

### D-alanylation of TAs is essential for the persistence of *Lactobacillus plantarum* in the *Drosophila* gut and the modulation of its immune-stimulatory potential as a core component of the gut-associated microbiota

In keeping with the role of D-alanine esterification of TAs in the resistance of *Bacilli* to antimicrobial effectors and the dampening of their IMD- eliciting potential, we asked whether this phenomenon would also account for the intestinal immune-tolerance of *L. plantarum* as a prevalent member of *Drosophila* microbiota. Therefore, we first determined whether the D-alanylation of TAs is required for the colonization and the persistence of *L. plantarum* in the *Drosophila* gut. The previously characterized *L. plantarum^WJL^* strain (*Lp^WJL^*), which was isolated from the intestine of *Drosophila melanogaster*, has a low transformation efficiency, which makes it difficult for site directed mutagenesis of the *dlt* operon. Therefore, we used the *L. plantarum^WCFS1^* strain (*Lp^WCFS1^*), isolated from human saliva, and its previously characterized *dlt* mutant (*Lp^WCFS1^Δdlt*)(Hayward & Davis, 1956) (Kleerebezem et al., 2003; Palumbo et al., 2006). In a recent study, Newell *et al*. have reported that the *Lp^WCFS1^* strain shares the same ability as the *Lp^WJL^* strain to colonize the *Drosophila* gut (Newell et al., 2014). These data were further confirmed in our hands (Figure 3 – figure supplement 1) thus; we checked whether *Lp^WCFS1^* reproduced the same *Lp^WJL^* triggered IMD-dependent immune response when introduced in the flies. For that, adult flies were fed on a mixture of antibiotics for three days to generate gut germ free flies (GF) (Stein, Roth, Vogelsang, & Nüsslein-Volhard), which were then recolonized with physiological quantities of the different strains of *L*. plantarum (Bonnay et al., 2013; Storelli et al., 2011). GF and recolonized states were attested by plating the dissected guts on appropriate agar media. The comparison of the *Diptericin* expression levels in the guts of flies conventionally reared on cornmeal agar medium (CR) to that of GF flies, shown in Figure 3 – figure supplement 1, indicates that the gut microbiota sets an IMD- dependent immune response. This immune response is mildly but significantly induced in guts recolonized with *Lp^WJL^* and *Lp^WCFS1^* as compared to GF guts. These results are in line with data previously reported by Bosco-Drayon *et al*. showing that *Lp^WJL^* is a mild inducer of AMP genes in the gut. These authors also reported that *Lp^WJL^* most potently activates the expression of genes encoding negative regulators of the IMD pathway such as *pirk* and *PGRP*-SC1. These results were also confirmed for the *Lp^WCFS1^* strain (Figure 3 – figure supplement 1). Collectively, our results attest that *Lp^WCFS1^* reproduces the IMD-dependent immune response activated by the *Drosophila* native gut associated *Lp^WJL^* strain. Therefore, we pursued our analysis by checking whether the *dlt* operon is required for the persistence of *L. plantarum* in the *Drosophila* gut. According to recent data, the *Drosophila* microbial community is sustained by continuous replenishment through feeding (Blum et al., 2013). Likewise, work from Leulier’s laboratory have previously shown that upon association in gnotobiotic flies, *L. plantarum* colonizes the *Drosophila* midgut and its external culture medium and resists the passage through the digestive tract of its host (Storelli et al., 2011). Therefore, in order to evaluate the persistence of the wt and *dlt* mutant strains in the *Drosophila* gut, GF adult flies were monoassociated by feeding on *Lp^WCFS1^* or the *Lp^WCFS1^*Δ*dlt* mutant strains for three days and then transferred to fresh cornmeal agar medium vials. The bacterial loads in the fly guts were assessed on daily basis. As shown in Figure 3A, the bacterial count remained invariable in the guts of wt flies mono-associated with Lp*^WCFS1^* whereas it significantly declined in the guts of the flies recolonized by the Lp*^WCFS1^Δdlt* mutant. In contrast, the bacterial loads of both wt and *Lp^WCFS1^Δdlt* mutants remained invariable in the guts of mono-associated *relish* mutants (Figure 3A). These results suggest that the D-alanylation of TAs is required for the resistance of *L. plantarum* to the IMD-dependent immune response and its subsequent persistence in the *Drosophila* gut. To further check whether this persistence particularly stems from the *dlt*-dependent resistance of *L. plantarum* to cationic antimicrobial effectors, we examined the persistence of the wt and *dlt* mutant strains of *Lp^WCFS1^* in the guts of transgenic flies overexpressing *Cecropin* (Figure 3 – figure supplement 2). As shown in Figure 3A, the counts of the wt *Lp^WCFS1^* strain were constant in the guts of the recolonized flies despite the overexpression of Cecropin, whereas those of the *Lp^WCFS1Δdlt^* mutant strain dramatically declined in less than four days. Altogether, these data underline a role of the *dlt* operon in the resistance of *Lp^WCSF1^* to cationic antimicrobial effectors thus favoring its persistence in the *Drosophila* gut.

**Figure 3:**
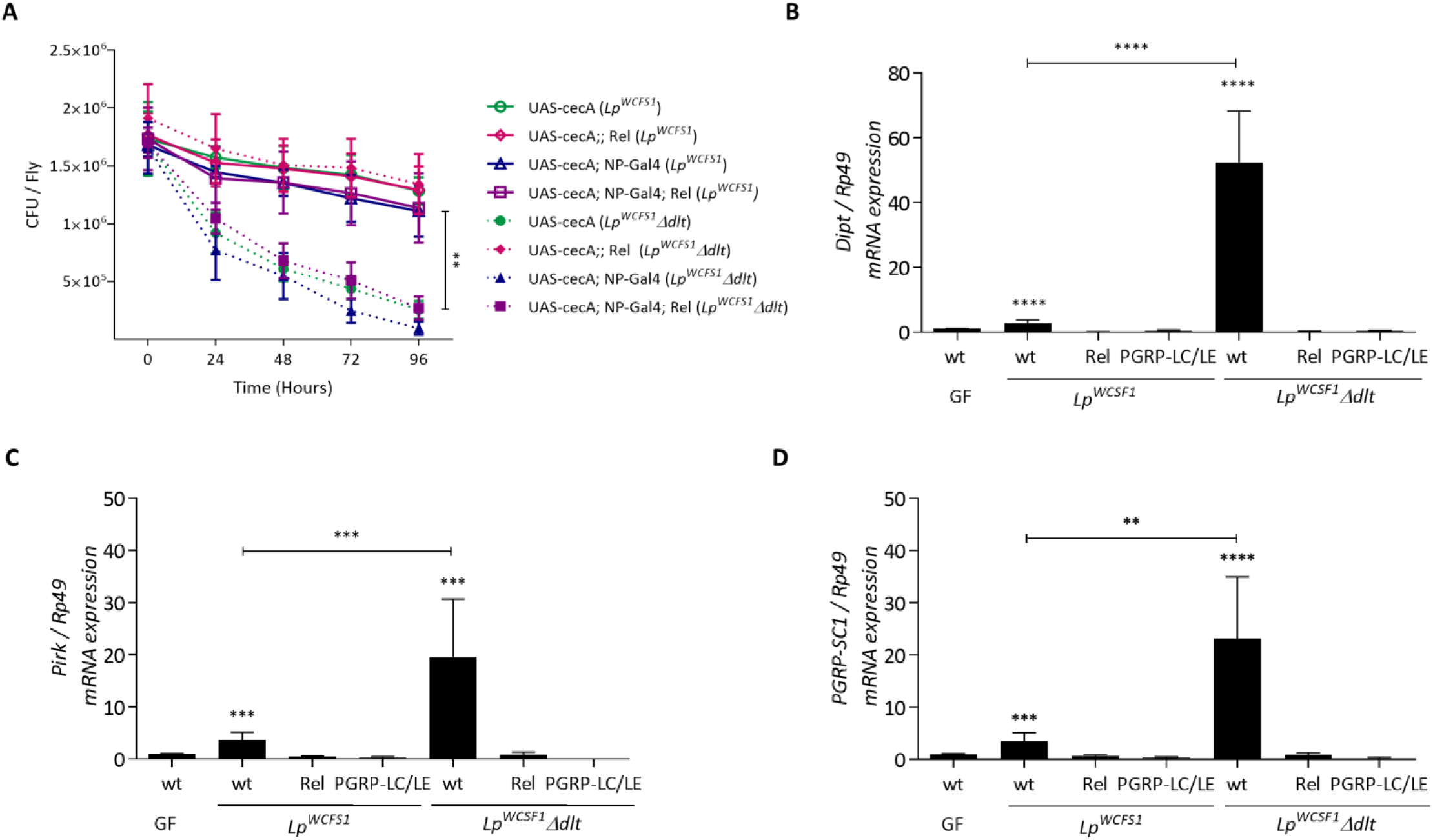
The D-alanylation of TAs is essential for the persistence and the immunomodulatory potential of *Lactobacillus plantarum* in the *Drosophila* gut. (A) Internal bacterial loads retrieved from the guts of Germ Free (GF) wild-type (wt) or *relish* (Rel) mutant flies fed with *Lp^WCFS1^* or *Lp^WCFS1^Δdlt* for three days. The expression of a *Cecropin* transgene is specifically driven in the gut of these flies by the NP-Gal4 driver. Colony Forming Unit (CFU) counting was performed every 24 hours after transferring the monoassociated flies to a fresh cornmeal medium vial. (B, C and D) Relative expression of the *Diptericin* (*Dipt*) (B) *Pirk*) (C) and *PGRP-SC1* (D) transcripts in the midgut of Germ Free (GF) wild-type (wt), *relish* (*Rel*) or *PGRP-LC/LE* double mutants flies fed with *Lp^WCFS1^* or *Lp^WCFS1^Δdlt* for 3 days. Transcripts expression was measured by RT-qPCR in total RNA extracts. Ribosomal protein 49 (Rp49) transcript was used as reference gene. Transcripts levels detected in GF flies is set as a control. Data obtained from three independent experiments are combined in single value (mean ± sd). Statistical tests were performed using the Mann-Whitney test within Prism software (ns: *p* > 0.05; *: 0.01< *p* < 0.05; **: 0.001< *p* < 0.01; ***: 0.0001< *p* < 0.001; ****: *p* < 0.0001). The exact *p* values are listed in Figure 3 – source data 1.

Next, we questioned whether the D-alanylation of TAs influenced the immune-stimulatory potential of *L. plantarum* in the intestine. To this end, we compared the intensity of the IMD response elicited in the guts of GF flies monoassociated with either *Lp^WCFS1^* or *Lp^WCFS1^Δdlt*. As shown in Figure 3B, the *Lp^WCFS1^Δdlt* mutant triggers a significantly accentuated immune response in the *Drosophila* gut as compared to that induced by the wt *Lp^WCSF1^* strain. Remarkably, the *Diptericin* expression level activated upon the sensing of *Lp^WCFS1^Δdlt* exceeds that of the negative regulators encoding genes (Figure 3C and 3D). The expression of all induced genes is strictly dependent on the activation of the IMD pathway as attested by the phenotype of *relish* or the *PGRP-LC/LE* double mutants (Figure 3B, 3C and 3D). These results contrast with those obtained for the wt *Lp^WCFS1^* strain. Of note, flies were daily transferred to new vials supplemented by the corresponding bacterial strains and variations in the ingested bacterial loads were excluded by the comparison of the CFU counts in the guts of flies monoassociated with the wt *Lp^WCFS1^*or the *Lp^WCFS1^Δdlt* mutant strains (Figure 3 – figure supplement 3). Taken together, our data indicate that the D-alanylation of TAs in *L. plantarum* is essential for the immune-modulatory potential of *L. plantarum* and its persistence in the *Drosophila* gut as a prevalent element of its microbiota.

### D-alanylation of TAs limits the release of PGN fragments through resistance to antimicrobial effectors

Our findings so far demonstrate a dual property for D-alanylation of TAs in *Bacilli*: first, this modification of the bacterial surface allows resistance to cationic antimicrobial effectors and second, it hampers sensing by the IMD pathway thus partially eluding the innate immune response. Previous studies have established that monomeric and polymeric fragments of PGN elicit the IMD immune response (Kaneko et al., 2004; Stenbak et al., 2004). Consistently, we reasoned that the *dlt*-dependent resistance to cationic antimicrobial effectors of *Bt407* and *L. plantarum* would limit any discharge of cell wall PGN thus only triggering a basal IMD response. To directly examine this possibility, we cultivated wt and *dlt* mutants of *Bt407* and *L. plantarum* bacterial strains in culture medium supplemented or not by chicken egg lysozyme. We then compared the IMD- induced response in adult flies injected with the supernatant of each of these microbial cultures. Of note, similarly to AMPs, lysozyme is a cationic antibacterial effector that drastically affects the growth of the *dlt* mutants in contrast to the highly resistant wt strains (data not shown). As shown in Figure 4, *Diptericin* expression triggered by the injection of the supernatants of *Bt407* and *Lp^WCFS1^* did not vary whether these cultures were treated or not with lysozyme (Figure 4A and 4B). In contrast, *Diptericin* expression significantly increased in flies injected with the supernatants of *Bt407Δdlt* and *LpWCFS1Δdlt* treated with lysozyme (Figure 4A and 4B). This response is strictly dependent on the activation of the IMD pathway as it is reduced in the *relish* mutant. Remarkably, the supernatants of the *dlt* cultures also elicited a significantly higher *Diptericin* expression as compared to that of wt bacteria. Collectively, our results suggest an increased discharge of immune-stimulatory PGN moieties in the supernatant of *dlt* mutants as compared to that of wt cultures.

**Figure 4:**
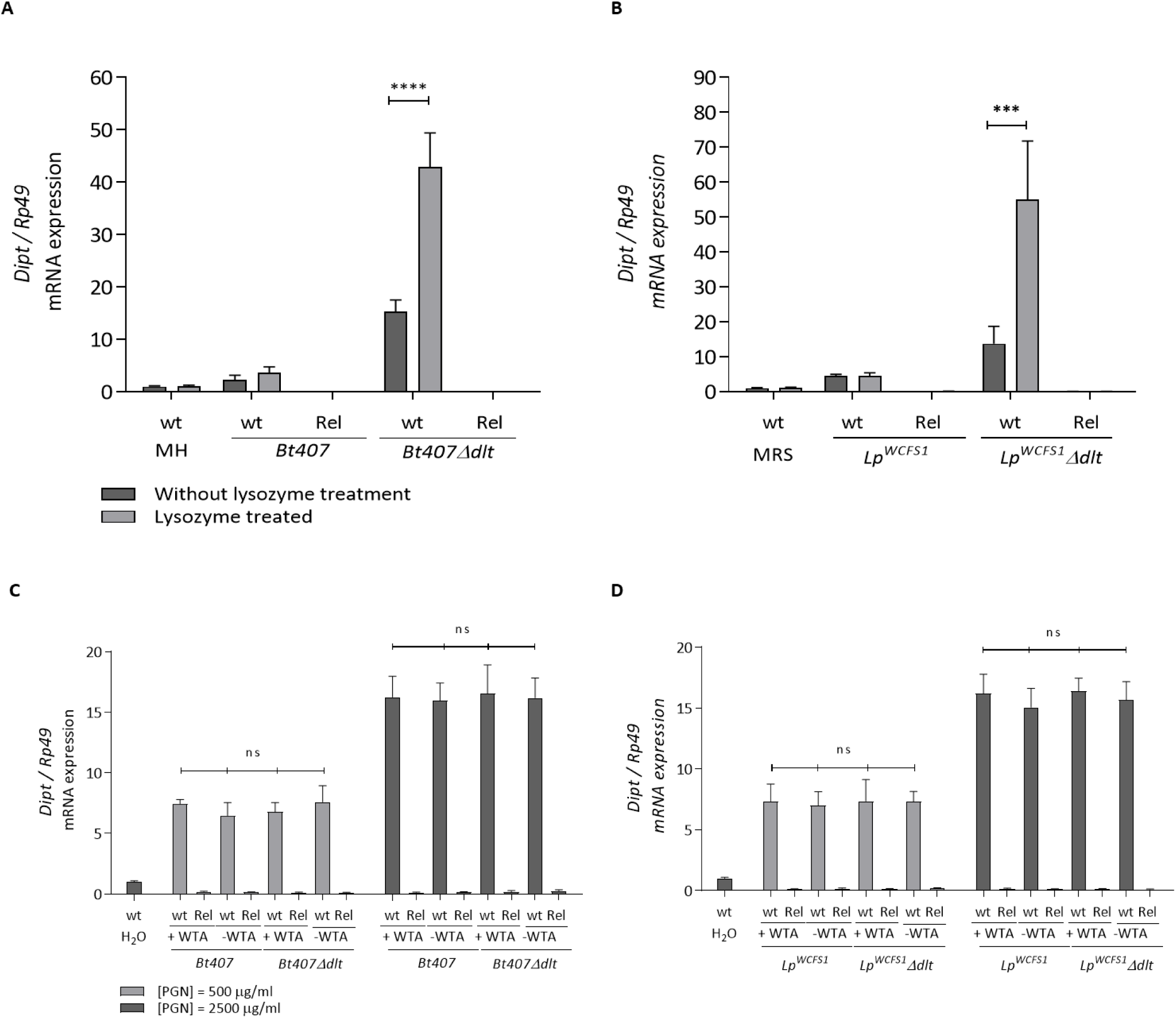
D-alanylation ot TAs limits the release of PGN fragments from the cell wall of *Bacilli* treated by cationic lysozyme. (A and B) Wild-type (wt) and *relish* (Rel) mutant flies were injected with the supernatant of *Bt407* and *Lp^WCSF1^* cultures or their corresponding *dlt* mutants (*Bt407Δdlt* and *Lp^WCSF1^Δdlt)* treated or not by chicken egg lysozyme (10 mg/ml). (C and D) Wild-type (wt) and *relish* (Rel) mutant flies were injected with cell wall peptidoglycan (PGN) fractions with or without cell wall TAs (500 and 2500 μg/ml) from the wild type or the *dlt* mutants of*Bt407* and *Lp^WCSF1^*. Relative *Diptericin* (dipt) expression was measured by RT-qPCR 4 h after the microbial induction of the adult flies. Ribosomal protein 49 (Rp49) transcript was used as reference gene. Transcripts levels detected in wt flies injected with MH Broth medium (A), MRS Broth Medium (B) or water (C and D) are set as controls. Data obtained from three independent experiments are combined in single value (mean ± sd). Statistical tests were performed using the Mann-Whitney test within Prism software (ns: *p* > 0.05; *: 0.01< *p* < 0.05; **: 0.001< *p* < 0.01; ***: 0.0001< *p* < 0.001; ****: *p* < 0.0001). The exact p values are listed in Figure 4 – source data 1.

Next, we asked whether the observed variations in the IMD-elicited response are indeed due to variable concentrations of the PGN fragments and not to any binding constraints related to the covalent bonding of D-alanylated TAs to the PGN fragments. Therefore, we purified cell wall fractions from the wt and *dlt* mutant bacteria and compared the IMD induced response upon their injection in adult flies. The extraction procedure was performed according to two different protocols in order to conserve or to eliminate TAs from the purified PGN fractions (see Experimental procedures). As shown in Figures 4C and 4D the *Diptericin* expression elicited upon the injection of cell wall fractions in the adult flies remains invariable whether these fractions were extracted from wt or *dlt* mutant strains. Moreover, this expression was not altered whether the TAs were removed from the PGN fractions or not. Interestingly, the elicited immune response also increased in a dose dependent manner, further supporting a direct correlation between the dose of bacterial PGN and the extent of the immune response. Thus, altogether, these data confirm that variations in the IMD-induced response elicited by wt and *dlt* mutants can be linked to varying doses of discharged PGN fragments.

### The *dlt*- dependent resistance of *Lactobacillus plantarum* to lysozyme is essential for modulating the IMD-dependent immune response in the *Drosophila* gut

Our results suggest that the D-alanylation of TAs mitigates the induction of the *Drosophila* IMD pathway by *B. thuringiensis* and *L. plantarum*. More precisely, by providing resistance to cationic antimicrobial effectors, D-alanine esterification of TAs constrains the release of PGN fragments from the bacterial cell wall. To evaluate the relevance of this model, we asked whether the depletion of antimicrobial effectors would alleviate the immune response triggered by the *dlt* mutants *in vivo*. However, as the *Drosophila* genome encodes several AMP genes and that the concomitant inhibition of their expression is quite difficult, we could not address this question in the frame of the systemic immune response triggered by a *B. thuringiensis* septic infection in flies. Therefore, we examined this possibility in the *Drosophila* gut where, in addition to AMPs, lysozyme constitutes another efficient cationic antimicrobial effector. The role of lysozyme in the activation of the IMD-response in the gut has not been considered to date. The *Drosophila* genome comprises seven lysozyme genes of which the members of the LysD family, including *LysD*, *Lys B*, *Lys C* and *Lys E*, and *LysX*, are expressed in the adult gut (Broderick et al., 2014; Daffre et al., 1994). As no lysozyme mutants are available, we expressed a dsRNA targeting the lysozyme transcripts under the control of the gut specific NP-gal4 driver (Hayashi et al., 2002). Since the lysozyme genes share high sequence similarities, the dsRNA constructs targeting the *LysS* and *LysX* transcripts also reduced the expression of all *LysD* like genes as off-targets. We further aggravated this RNAi transcript-depletion by introducing a genetic deficiency uncovering the lysozyme locus into the genome of the transgenic flies (Figure 5 – figure supplement 1). Since we could not establish an oral infection of flies with *B. thuringiensis*, we evaluated the role of gut lysozyme in the modulation of the immune response triggered by the gut resident *L. plantarum*. For that, we quantified *Diptericin* expression level in the guts of GF- RNAi flies recolonized by either the *Lp^WCFS1^* or the *Lp^WCFS1^Δdlt* mutant strains. As shown in figure 5, compared to control RNAi-GFP flies, the inhibition of lysozyme genes significantly reduced expression of *Diptericin* in flies mono-associated with *Lp^WCFS1^Δdlt* and the level of reduction in the lysozyme transcripts correlated with the level of reduction in *Diptericin* expression. Of note, *Diptericin* expression did not vary in RNAi treated flies replenished by *Lp^WCFS1^* whether lysozyme expression was inhibited or not. These results are in agreement with an increased discharge of PGN fragments from *Lp^WCFS1^Δdlt* as a result of its vulnerability to cationic antimicrobial effectors *in vivo* and provide a first evidence of the role of lysozyme in modulating the IMD response in the *Drosophila* gut.

**Figure 5:**
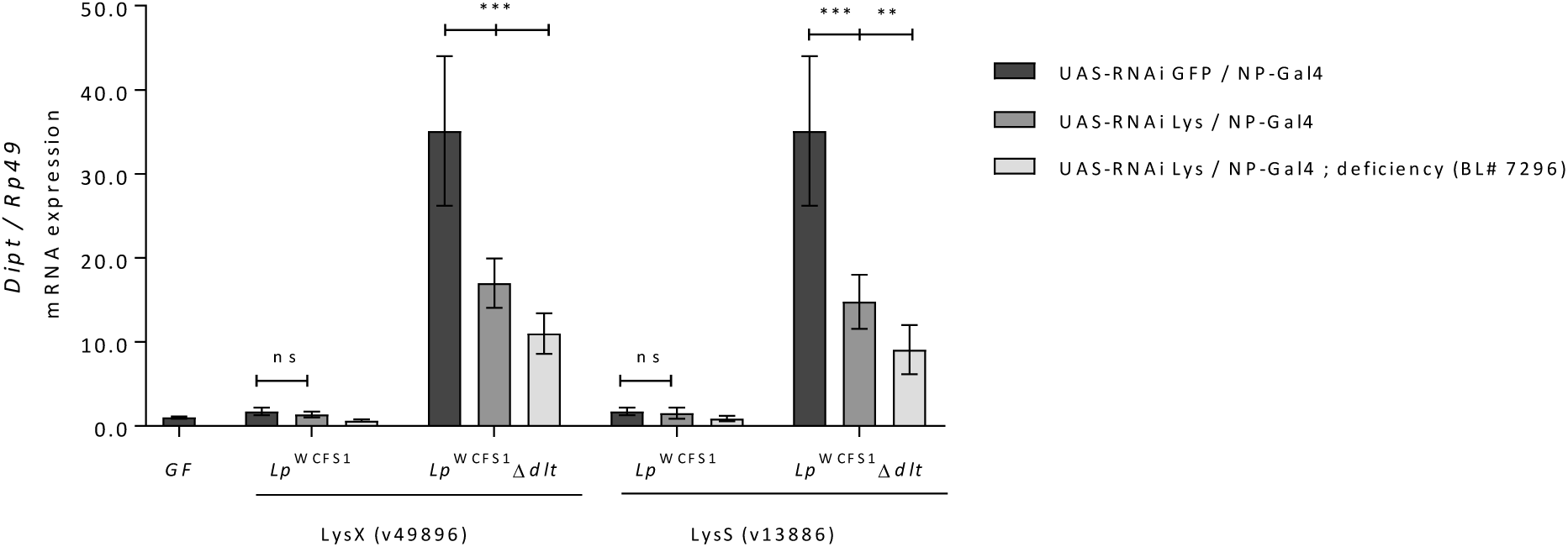
The *dlt*-dependent resistance of *Lactobacillus plantarum* to intestinal lysozyme is essential for the modulation of its ensued IMD- immune response in the gut. Silencing of the lysozyme genes was performed by the expression of RNAi constructs (UAS-RNAi) in the gut of adult flies using the midgut specific driver NP-Gal4. Two different transgenic fly lines (v49896) and (v13886) were used. These harbor different RNAi construct targeting the *LysX* or the *LysS* transcripts respectively. Silencing of *lysozyme* genes is further aggravated by crossing these transgenic flies to a fly line carrying a chromosomal deletion (deficiency BL#7296) uncovering the *lysozyme* locus in the *Drosophila* genome. A transgenic line expressing an RNAi construct targeting the GFP transcript is used as a control. Relative expression of the *Diptericin* transcript is measured in total RNA extracts from the midgut of Germ Free wild-type flies (GF) or the transgenic flies of the different genetic background fed on *Lp^WCFS1^* or *Lp^WCFS1^Δdlt* for 3 days. Ribosomal protein 49 (Rp49) transcript was used as reference gene. Transcripts levels are compared to that detected in in GF flies that is arbitrary set to 1. Data obtained from three independent experiments are combined in single value (mean ± sd). Statistical tests were performed using the Mann-Whitney test within Prism software (ns: *p* > 0.05; *: 0.01< *p* < 0.05; **: 0.001< *p* < 0.01; ***: 0.0001< *p* < 0.001; ****: *p* < 0.0001). The exact p values are listed in Figure 5 – source data 1.

## Discussion

The role of the *dlt*-operon in mediating resistance to cationic antimicrobial effectors *in vitro* has been attested for several Gram-positive bacteria species (Fabretti et al., 2006; Kovacs et al., 2006; Kristian et al., 2005; Perea Velez et al., 2007). This attribute is associated with an attenuated virulence of the *dlt* mutants for several pathogens including *Listeria monocytogenes*, *Bacillus anthracis* and *Staphylococcus aureus* (Abachin et al., 2002; Collins et al., 2002; Fisher et al., 2006; Kristian et al., 2003; Tabuchi et al., 2010). Only for the latter, a role for the *dlt*-operon in hampering immune sensing and the subsequent activation of the Toll pathway in *Drosophila* has been proposed (Tabuchi et al., 2010). The Toll pathway is preferentially triggered upon the sensing of Lys-type PGN by a receptor complex comprising PGRP-SA and GNBP1, a member of the *Drosophila* Glucan-Binding protein family (Gobert et al., 2003; Michel, Reichhart, Hoffmann, & Royet, 2001). However, although the exact mechanism underlying the inhibitory effect of D-alanine esterification of TAs on the Toll pathway activation remains unknown, the results obtained by Kurokawa *et al*. suggest that it entails the covalent bonding of D-alanylated Wall TAs to PGN (Kurokawa et al., 2011). This model is challenged by a report from Atilano *et al*. which indicates that wall TAs limit the access of PGN to the PGRP-SA receptor independently from the activity of the *dlt*-operon (Atilano, Yates, Glittenberg, Filipe, & Ligoxygakis, 2011). We show here that, the presence of TAs, whether D-alanylated or not, does not alter the IMD response upon injection of purified PGN preparations in flies (Figure 4B). Furthermore, the quantification of the IMD response in flies stimulated by injection of supernatants of bacterial cultures treated or not by chicken egg lysozyme, or increasing concentration of PGN preparations, reveals a correlation between the induced response and the doses of injected PGN fragments (Figure 4 A and B). Overall, our data support a model in which the differential immune response following infection of flies by wt or *dlt* mutant bacteria is linked to variations in the quantities of released PGN that are made available for sensing by the PGRP-LC/ LE receptors upon lysis by AMPs. The differences between our results and those documented for Toll pathway activation might be indicative of different ligand binding / receptor activation mechanisms for the PGRP-LC/ LE and the PGRP-SA/GNBP1 PRR complex.

Resistance to antimicrobial effectors and evasion from immune detection are prominent features of pathogenic bacteria. Interestingly, our data indicate that the D-alanylation of TAs is a common microbial strategy employed by pathogen and commensal alike to elude and resist immune defenses thus acquiring the capacity to develop or persist in appropriate niches. Indeed, our results reveal the essential role of D-alanine esterification in the resistance of *L. plantarum* to the assault of intestinal cationic lysozyme and thus for its persistence as a core component of the *Drosophila* microbiota (Figure 3A). These findings are in agreement with a previous report attesting of the prominence of D-alanine esterification of TAs for the colonization of *L. reuteri* in the mouse intestinal tract (Perea Velez et al., 2007; Walter et al., 2007). The *dlt* operon was also found to be an essential element for the establishment of *L. casei* in the gut of the rabbit ligated ileal loop model (Licandro-Seraut, Scornec, Pedron, Cavin, & Sansonetti, 2014). Our data also point the crucial role of the D-alanylation of TAs in maintaining the immunomodulatory potential of *L. plantarum* in the *Drosophila* gut. This cell wall modification allows the fine-tuning of IMD signaling as to set an essential immune barrier in the gut, which is a prerequisite for the maintenance of a healthy commensal population. Indeed, several lines of evidence have indicated the harmful effects of an exacerbated IMD response in the *Drosophila* gut (Bonnay et al., 2013; Lhocine et al., 2008; Paredes et al., 2011; Ryu et al., 2008). Notably, a dysregulated IMD response has been associated with commensal dysbiosis and epithelial dysplasia (Guo et al., 2014; Ryu et al., 2008). These data further highlight the importance of the dual role of D-alanylation of TAs in the resistance to antimicrobial effectors and the escape of immune detection as defining characteristics of *L. plantarum* as a *bona fide* beneficial *Drosophila* commensal. In other words, resistance to antimicrobial effectors allows the persistence of *L. plantarum* in the gut and in the habitat of its host. Whereas limiting the discharge of its PGN immunostimulants, permits the set-up of a steady state immune response that would not perturb the equilibrium of gut commensals. Remarkably, according to our results, this immunomodulation capacity of *L. plantarum* largely relies on its *dlt*-dependent resistance to gut lysozyme (Figure 3B, 3C and 3D). Our genetically based data support the long surmised immunomodulatory role of lysozyme (Callewaert & Michiels, 2010; Ragland & Criss, 2017).

Immunomodulation is as an essential property by which lactic acid bacteria exert their biological functions in mammals. It is also the main criteria for the selection of probiotic strains (Bron, Tomita, Mercenier, & Kleerebezem, 2013; I. C. Lee, Tomita, Kleerebezem, & Bron, 2013; Wells, 2011). Notably, the cytokine profiles triggered upon the sensing of these bacteria in the intestinal mucosa determine the subsequent polarization of T cells, and thus the orientation of the ensuing adaptive immune response. Differential cytokine profiles are triggered upon the sensing of appropriate Microbial Associated Molecular Patterns (MAMPs) of *lactobacilli* by cognate PRRs at the intestinal epithelial barrier (Lebeer, Vanderleyden, & De Keersmaecker, 2010). Notably, previous data have suggested that the sensing of lipoteichoic acid (LTA) is mainly performed by the transmembrane TLR2 receptor, whereas sensing of bacterial PGN fragments is preferentially mediated by intracellular NOD receptors (Macho Fernandez, Pot, & Grangette, 2011; Smelt et al., 2013; Travassos et al., 2004). Interestingly, the wt and the *dlt*-mutant of the *Lp^WCFS1^* strain triggered a differential cytokine profiles in mammals (Grangette et al., 2005; Smelt et al., 2013). The transcriptional response triggered by the former is marked by the expression of TLR2-dependent pro-inflammatory IL-12 whereas that activated by the latter is characterized by an enhanced expression of anti-inflammatory IL-10 that is mainly NOD-dependent (Grangette et al., 2005; Macho Fernandez, Pot, et al., 2011; Macho Fernandez, Valenti, et al., 2011). Similar results were obtained with a wt and *dlt*-mutant strain of *L. rhamnosus GG* (Claes et al., 2010). Accordingly, the *dlt* mutants alleviated the inflammatory response in a mouse colitis model (Claes et al., 2010; Grangette et al., 2005). Despite the complexity of the immune response in the mammalian intestinal mucosa, these data are reminiscent of our results reporting a differential immune response triggered by the wt and the *Lp^WCFS1Δdlt^*. Interestingly, a recent study from Bosco-Drayon and colleagues emphasized the prominent role of the intracellular PGRP-LE receptor as a master sensor of small fragments of PGN (tracheal cytotoxin) in the *Drosophila* gut further underlying the similarity between the function of this receptor and that of mammalian NODs (Bosco-Drayon et al., 2012). Given the sensitivity of the *dlt* mutants of Lactobacilli to antimicrobial effectors (Perea Velez et al., 2007; Walter et al., 2007) and the high occurrence of lysozyme and AMPs at the mammalian intestinal barrier, it is tempting to speculate that a similar mechanism might apply for the regulation of PGN-sensing and the modulation of innate immune responses by commensal and probiotic *Lactobacilli* species in *Drosophila* and mammals (Lebeer et al., 2010; Perea Velez et al., 2007).

Several reports have demonstrated the potential of *L. plantarum* to impact *Drosophila* development and physiology (Blum et al., 2013; Bosco-Drayon et al., 2012; Broderick et al., 2014; Erkosar & Leulier, 2014; Guo et al., 2014; Jones et al., 2013; Matos et al., 2017; Ryu et al., 2008; Sharon et al., 2010; Storelli et al., 2011; A. C. Wong et al., 2014). Notably, work from Leulier’s laboratory revealed the importance of this bacterium in promoting juvenile growth during chronic undernutrition (Storelli et al., 2011). Interestingly, Matos *et al*. showed that a functional *dlt*-operon is required for this biological function of *L. plantarum* (Matos et al., 2017). Our data give further evidence of the importance of the *dlt*-dependant D-alanylation of TAs for the beneficial impacts of *L. plantarum* on its host biology in particular by tuning the sensing of PGN immunostimulants, thus establishing an *ad hoc* balanced immune response at the intestinal epithelium. Moreover, in *Drosophila* as in mammals, *L. plantarum* was shown to exert protective probiotic effects. Notably, in a recent study, Blum *et al*. provided evidence of a protective role of *L. plantarum* against intestinal pathogens such as *Serratia marcescens*. In this context, complementation of fly food by *L. plantarum* significantly enhanced the capacity of CR and GF flies to survive an infection by *S. marcescens* (Blum et al., 2013). Interestingly, we showed that this probiotic effect of *L. plantarum* feeding in *Drosophila* is strictly dependent on the activity of its *dlt* operon as evidenced by the phenotype of flies fed by the *Lp^WCFS1Δdlt^* mutant strain (Figure 3 – figure supplement 4). These data are in line with a recent comprehensive study performed by Smelt *et al*., which demonstrated that the majority of the immunomodulatory effects induced by the probiotic *Lp^WCFS1^* strain in mice are dependent on the activity of its *dlt*-operon (Smelt et al., 2013).

To summarize, our results put forward the pivotal role of the D-alanine esterification of TAs in modulating the sensing of bacterial PGN by the host immune system by preventing antimicrobial effectors mediated bacterial lysis. Modification of bacterial PGN was previously reported as an immune evasion strategy of pathogens to prevent its degradation by lysozyme and thus the release of immunostimulatory fragments (Boneca et al., 2007; Wolfert, Roychowdhury, & Boons, 2007). The function of the *dlt*-operon is unique in the sense that the modification of a cell-wall component, namely TAs, interferes with the sensing of another component, which is PGN, by the host innate immune sensors. Although the *dlt* operon is highly conserved among Gram-positive species, including saprophytic bacteria, such as the harmless and non-pathogenic *B. subtilis*, our results provide a clear evidence that its function is crucial for the success of pathogenic and commensal species in resisting host innate immune defenses. Given the high conservation of the innate immune system and the prevalence of TAs and PGN as central modulators of host defenses, these findings might yield new insights into the pathogenesis and the probiotic effects of bacterial species and potentially orient the development of therapeutic and prophylactic interventions.

## Materials and Methods

### Bacterial strains and growth conditions

The *dlt* operon mutant of *B. thuringiensis* was generated from the strain *B. thuringiensis* 407 cry-strain (*Bt407*) (Lereclus, Arantes, Chaufaux, & Lecadet, 1989) as described in (Abi Khattar et al., 2009). Briefly, the demethylated pMAD-updlt-aphA3Km-downdlt plasmid (Abi Khattar et al., 2009) was prepared from *Escherichia coli* ET12567 and introduced, by electroporation, into *Bt407*. The *dltXABCD* genes were replaced with the kanamycin (km) resistant cassette by allelic exchange. Clones resistant to Km and susceptible to Erythromycin (Em) were examined by PCR, and the homologous recombination was verified by sequencing.

The *dltA* mutant of *B. subtilis* was constructed in strain 168 as follows: an internal fragment of the *dltA* gene was amplified with primers BSdltA-*Eco*RI (5’-ggaattcctcggccagcagattcagacagttt-3’) and BSdltA-BamH1 (5’-acccggatccatcaggcacatttg-3’). The PCR product and the integrative plasmid pDIA5307 (PMID: 8113162) were digested with *Eco*RI and *Bam*HI and ligated together generating pDA5307-BSdltA. The ligated product was digested with *Sma*I to linearize any pDIA5307 plasmid left, transformed into competent cells of strain 168 and selected on chloramphenicol (4 µg/ml). Clones were validated by PCR and 4 independent clones were stored generating strain 168 Δ*dltA*. Clone 1 was used in this work.

*Serratia marcescens* Db11 (Nehme et al., 2007) and *E. coli* DH5αGFP (Bonnay et al., 2013) are a kind gift from Dr. Dominique Ferrandon and Pr. Jean Marc Reichhart (UPR9022 – CNRS - Université de Strasbourg) and *L. plantarum* WCFS1 (*Lp ^WCFS1^*) (Kleerebezem et al., 2003) and its *Δdlt* mutant (*Lp ^WCFS1^Δdlt*) (Palumbo et al., 2006) from Pr. Pascal Hols (Unité de Génétique, Institut des Sciences de la Vie, Université catholique de Louvain, B-1348 Louvain-La-Neuve, Belgium). All bacterial strains, except *L. plantarum*, were cultured on Luria-Bertani medium (LB) at 37°C and 30°C for *B. thuringiensis*. *L. plantarum* was cultured on Man, Rogosa and Sharpe (MRS) medium at 37°C.

Heat killing of bacteria was performed as described (El Chamy, Leclerc, Caldelari, & Reichhart, 2008) Briefly, bacterial solutions followed two steps of 20 minutes of incubation at 95°C separated by 20 minutes of cooling on ice. Killing was verified by plating 100 μl of each bacterial solution on LB agar plates.

For the preparation of the supernatant of bacterial cultures treated with Lysozyme, chicken egg white lysozyme (10mg/ml) was applied to bacterial cultures of *Lp^WCFS1^, Lp ^WCFS1^Δdlt, Bt407* and *Bt407Δdlt* at an optical density of 0.5 at λ = 600 nm (OD_600_ = 0.5) in Muller Hinton (MH) and MRS mediums respectively. When the OD_600_ reached 2, the bacterial cultures were centrifuged, and supernatants were heated at 95°C for 20 minutes before injection in the flies.

### Quantification of D-Alanine from TAs and purification of bacterial cell wall fractions

Quantification of D-Ala esterified to TAs was performed by HPLC as described previously (Kamar et al., 2017), after release by alkaline hydrolysis of whole cells or cell wall fractions. Cell walls were prepared from *Lp^WCFS1^, Lp ^WCFS1^Δdlt, Bt407* and *Bt407Δdlt* as described (Bernard et al., 2011). SDS, Pronase (2 mg/ml), Trypsin (200 µg/ml), DNase (50 µg/ml) and RNase (50 µg/ml) treatments were applied to obtain cell walls containing peptidoglycan and covalently attached glycopolymers (WTA, polysaccharides). All incubations were performed at pH ≤ 6.0, to keep D-Ala esterified on WTAs. The presence of D-Ala in the purified cell walls was checked by HPLC as described above.

Purified peptidoglycan was obtained after incubation of cell walls in 48% hydrofluoric acid at 4°C for 16 hours, to remove WTA and other polysaccharides attached covalently to peptidoglycan.

### Drosophila melanogaster stocks and maintenance

Oregon-R is used as wild-type. The IMD pathway mutants Relish^E20^ (*Rel*), *PGRP-LC^ΔE12^* (PGRP-LC) and the *PGRP-LC/LE* double mutant combines the *PGRP-LC^ΔE12^* and the *PGRP-LE^E112^* mutations were been described (Gottar et al., 2002; Hedengren et al., 1999; Takehana et al., 2004). The UAS-Cecropin A (UAS-Cec) (Reichhart et al., 2002) is a kind gift from Pr. Jean Marc Reichhart (UPR9022 – CNRS - Université de Strasbourg). Flies carrying an UAS-RNAi against *Lysozyme S* (v13386 GD), *Lysozyme X* (v49896 GD) were obtained from the Vienna *Drosophila* RNAi Center (http://stockcenter.vdrc.at/control/main). UAS-RNAi GFP line (397-05) were obtained from the *Drosophila* Genetic Resource Center (Kyoto, Japan; http://www.dgrc.kit.ac.jp/index.html). The deficiency Df(3L)10F125/TM6B,Tb (BL# 7296) and the fat body driver c564-Gal4 (BL# 6982) are from the Bloomington Stock Center. The adult gut tissues driver NP3080-Gal4 (NP-Gal4) was obtained from the *Drosophila* Genetic Resource at the National Institute of Genetics (Shizuoka, Japan). The Gal4 system activation was performed by incubating 2-3 days old flies for six further days at 29 °C. Fly stocks were raised at 25°C on cornmeal-agar medium rich in yeast (7.25%). For antibiotic treatment the medium was supplemented with a mixture of Ampicillin, Kanamycin, Tetracyclin and Erythromycin at 50 ug/ml each (final concentration). Adult female flies, aged between 0 to 1 day old, were fed on this mixture for 3 days. Germ free status was checked by plating adult fly gut on MRS and LB agar mediums.

### Infections and survival experiments

Survival experiment were performed on a total of ≥ 45 adult females per genotype (15 to 20 individuals per each of the three biological replicates). Batches of 15 to 20 female flies, aged between 2 to 4 days old, were pricked with a tungsten needle previously dipped into a bacterial solution prepared from an overnight culture that was washed and diluted in PBS (1x) to a final OD_600_ = 2. The infected flies were incubated at 29 °C and survival were counted were counted 15, 21 and 25 hours post infection. The *Diptericin* expression was measured by RT-qPCR 4h post infection.

Oral infection of flies by *S. marcescens* was performed as described (Nehme et al., 2007). In brief, an overnight bacterial culture was centrifuged and diluted to a final OD_600_ = 1 in a 50mM sucrose solution. The challenged flies were incubated on 29 °C for 54 h and the *Diptericin* expression was quantified in RNA extracts of 5 to 10 dissected guts by RT-qPCR.

9.2nl of sonicated purified peptidoglycan or 69 nl of supernatant treated with lysozyme were injected with into the thorax of batches of 20 to 25 female flies (aged 2 to 4 days old) with a Nanoject apparatus (Drummond, Broomall, PA). These flies were incubated 4 hours at 29 °C then *Diptericin* expression was quantified in RNA extracts by RT-qPCR.

### Fly internal bacterial load quantification

The CFU counting was performed on 30 adult females or 15 to 30 midguts per genotype (10 adults or 5 to 10 midguts per each of the three biological replicates) by plating serial dilutions of lysates obtained from infected flies or midguts on LB or MRS mediums containing the appropriate antibiotic to each strain. Each biological sample was analyzed by two technical replicates per experiment.

### Monoassociation of germ-free flies

Germ free female flies were transferred to a new food fly medium covered with 150 μl of bacterial solution prepared from an overnight culture of *Lp^WCFS1^* or *Lp ^WCFS1^Δdlt* washed and diluted in MRS to a final OD_600_ = 1. *Diptericin*, *pirk*, *PGRP-SC1* expressions were quantified by RT-qPCR from RNA extracts of 5 to 10 dissected guts from flies fed 3 days on *L. plantarum* feeding.

### Maintenance of *L. plantarum* in *Drosophila*’s gut

Batches of 20 Germ free flies were fed *Lp^WCFS1^* or *Lp ^WCFS1^Δdlt* for 3 days and then transferred to a fresh cornmeal medium vial. Every 24 hours, 5 to 10 guts were dissected and plated on MRS Medium containing the appropriate antibiotic to each strain.

### Quantification of gene expression

Total RNA was extracted with TRI Reagent (Sigma – Aldrich) from 15 to 25 adult females or 5 to 10 dissected adult female guts of each genotype per biological replicate. The reverse transcription was performed on 1 μg of RNA by using the RevertAid RT Reverse Transcription Kit (Thermo Fisher Scientific). cDNA was used as template in PCR reaction with three 10 μl technical replicates of each biological sample. qPCR was performed on an iQ5 Real Time PCR detection system (Biorad) using iTaq Universal Syber Green supermix (Biorad). The amount of RNA detected was normalized to that of the house keeping gene *rp49*. Primers for *Diptericin*, *Attacin D*, *pirk*, *PGRP-SC1*, *rp49* and *lysozymes* genes are listed in the supplementary file 1. Relative gene expression levels between control and experimental samples was determined using the ΔΔCT method. Each experimental sample was compared to each wild-type sample.

### Statistical analysis

Statistical analysis was performed using GraphPad Prism 7.0b software (α = 0.05) and statistical tests used for each data set are indicated in figure legends.

## Acknowledgments

We thank Pr. Jean-Marc Reichhart and Dr. Dominique Ferrandon for critical reading of the manuscript. We thank Pr. Pascals Hols for providing the *Lp ^WCFS1^* strain and its *Δdlt* mutant. This work was funded by the Research Council of the Saint-Joseph University of Beirut (FS73) and the Lebanese National Council of Scientific Research (CNRS-L/ 02-05-16) and has benefited from the support of the CEDRE program (32942VD) and Institut National de la Recherche Agronomique (INRA). Zaynoun Attieh is supported by a PhD fellowship from the CNRS-L and has also benefited from the SAFAR program and an international mobility scholarship from AgroParisTech (Bourse de mobilité Abies). *B. subtilisΔdlt* mutant was constructed in the frame of a European Research Council starting grant (PGNfromSHAPEtoVIR n°202283) awarded to Ivo G. Boneca.

## Figures Supplements

**Figure 1 – figure supplement 1:**
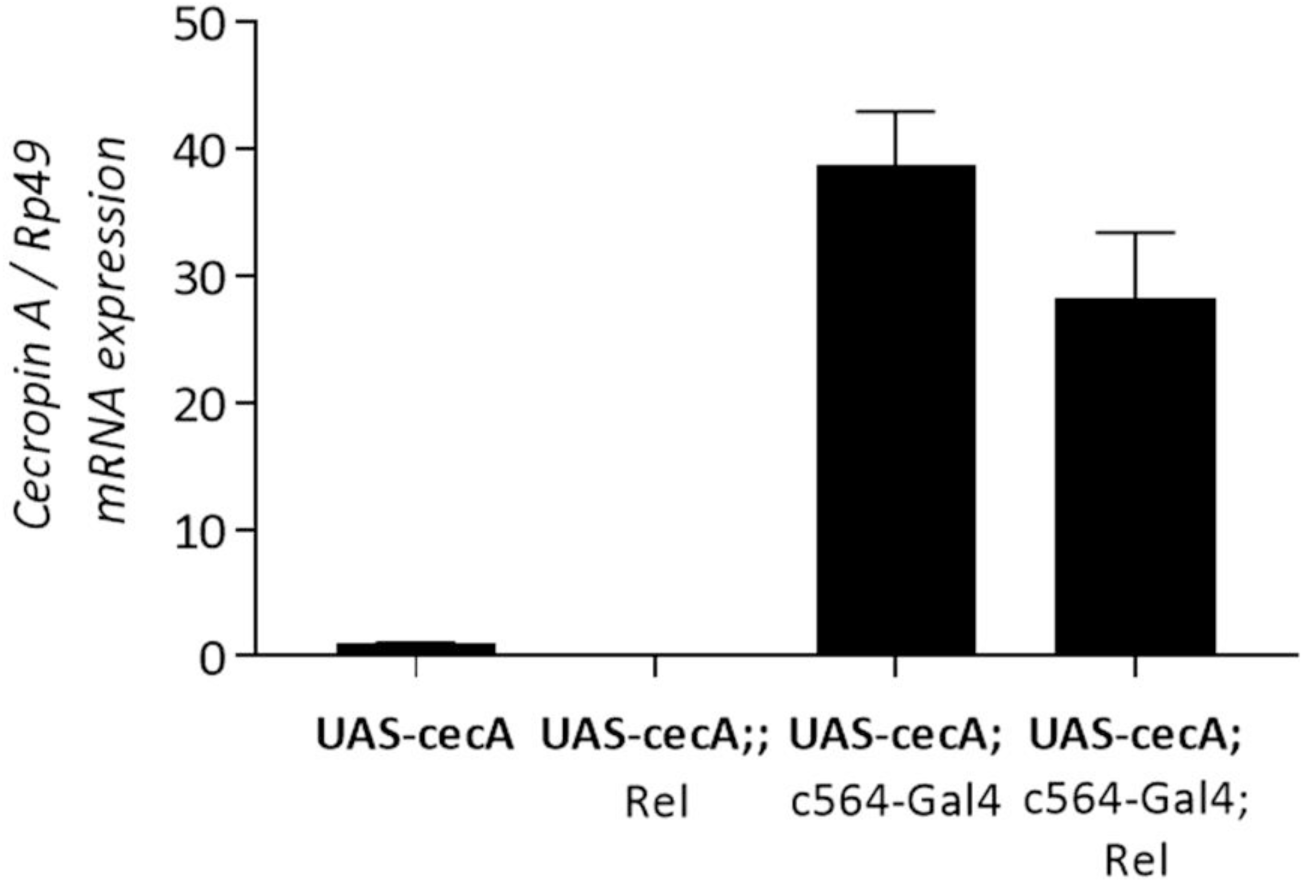
Overexpression of *CecropinA* gene. The expression of a *Cecropin A* transgene (UAS-cec) was driven using a fat body driver, c564-Gal4, in wild-type (wt) or *Rel* mutant flies. The *Cecropin A* expression a was quantified by RT-qPCR from the RNA extracts of adult flies following their incubation for 6 days at 29 °C. Wt flies carrying the UAS- cec transgene were used as control. Data obtained from three independent experiments are combined in single value (mean ± sd).

**Figure 1 – figure supplement 2:**
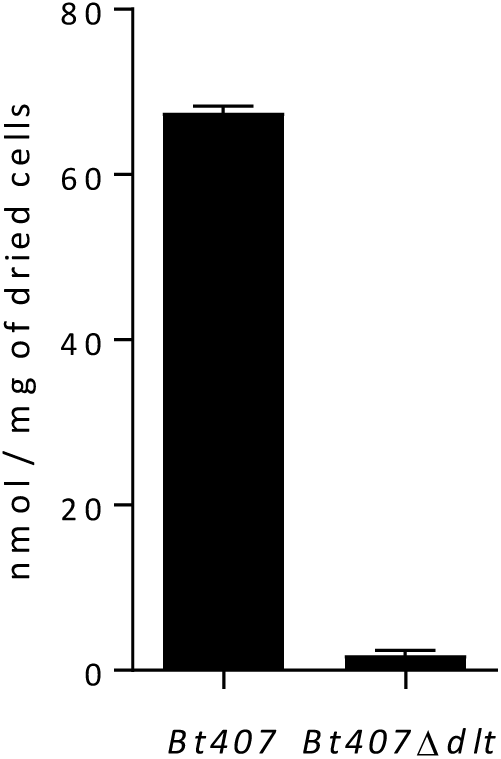
Amounts of D-alanine released from whole cells by alkaline hydrolysis for the wild type and the *dlt* mutant strains of *Bt407*. Data obtained from three independent experiments are combined in single value (mean ± sd).

**Figure 1 – figure supplement 3:**
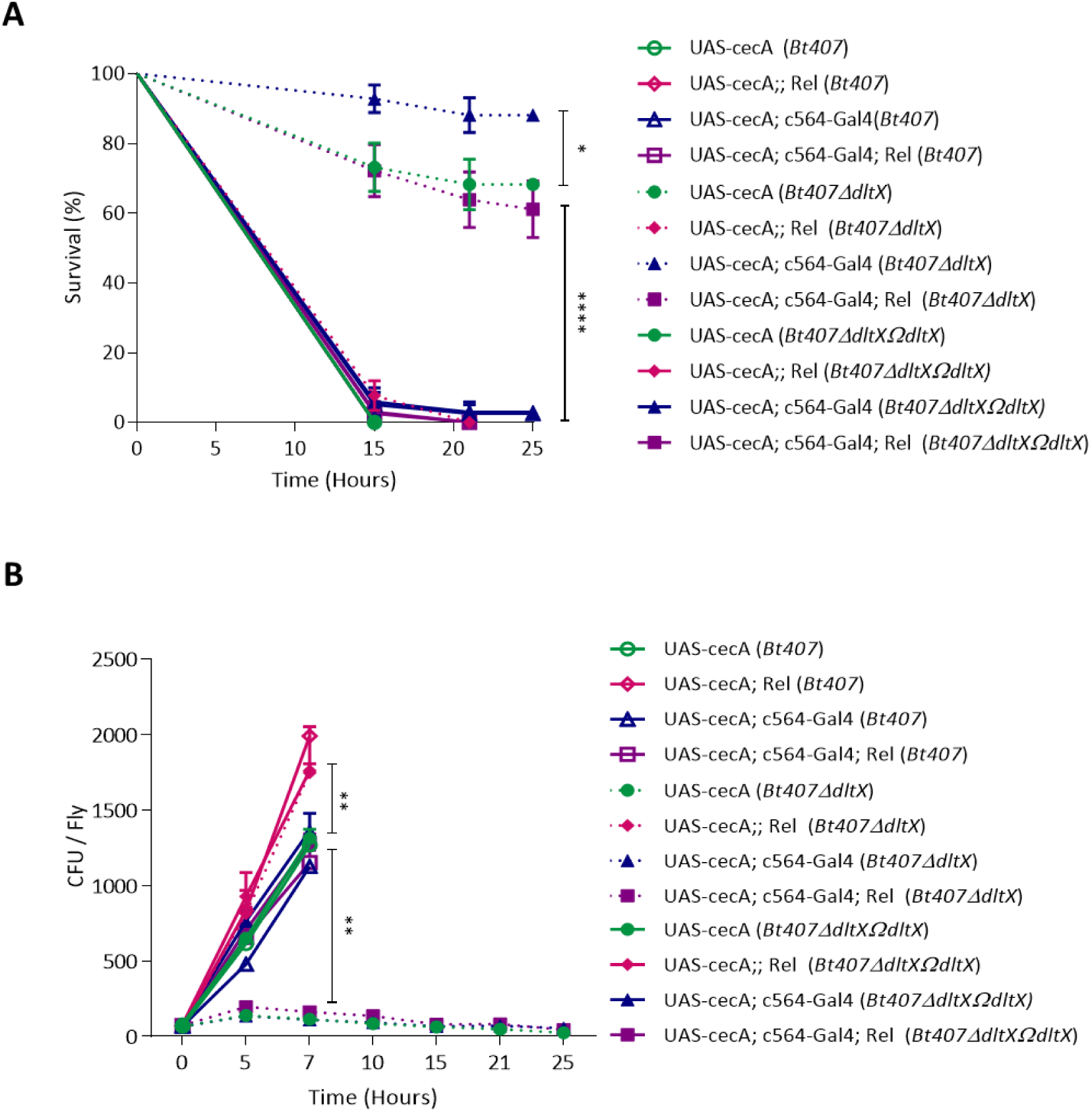
Complementation of the *Bt407ΔdltX* mutant with the *dltX-ORF* (*Bt407ΔdltΩdltX*) completely restores its virulence similarly to the wt *Bt407* strain. (A) Survival of adult wild-type (wt) or relish (Rel) mutant, overexpressing or not a *Cecropin* transgene (under the control of the c564-Gal4 fat body driver) to an infection with *Bt407, Bt407ΔdltX and Bt407ΔdltX ΩdltX* (B) Internal bacterial loads retrieved from adult flies infected with *BT407*and *Bt407ΔdltX* or *Bt407ΔdltX ΩdltX*. The Colony Forming Unit (CFU) counting was performed only on survival flies. Data are pooled from three independent experiments (mean and s.d.). Statistical tests were performed using the Log Rank test for the survival assays and Mann-Whitney test for the CFU counting within Prism software (ns: *p* > 0.05; * 0.01< *p* < 0.05; **:0.001< *p* < 0.01; ***: 0.0001< *p* < 0.001; ****: *p* < 0.0001).

**Figure 3 – figure supplement 1:**
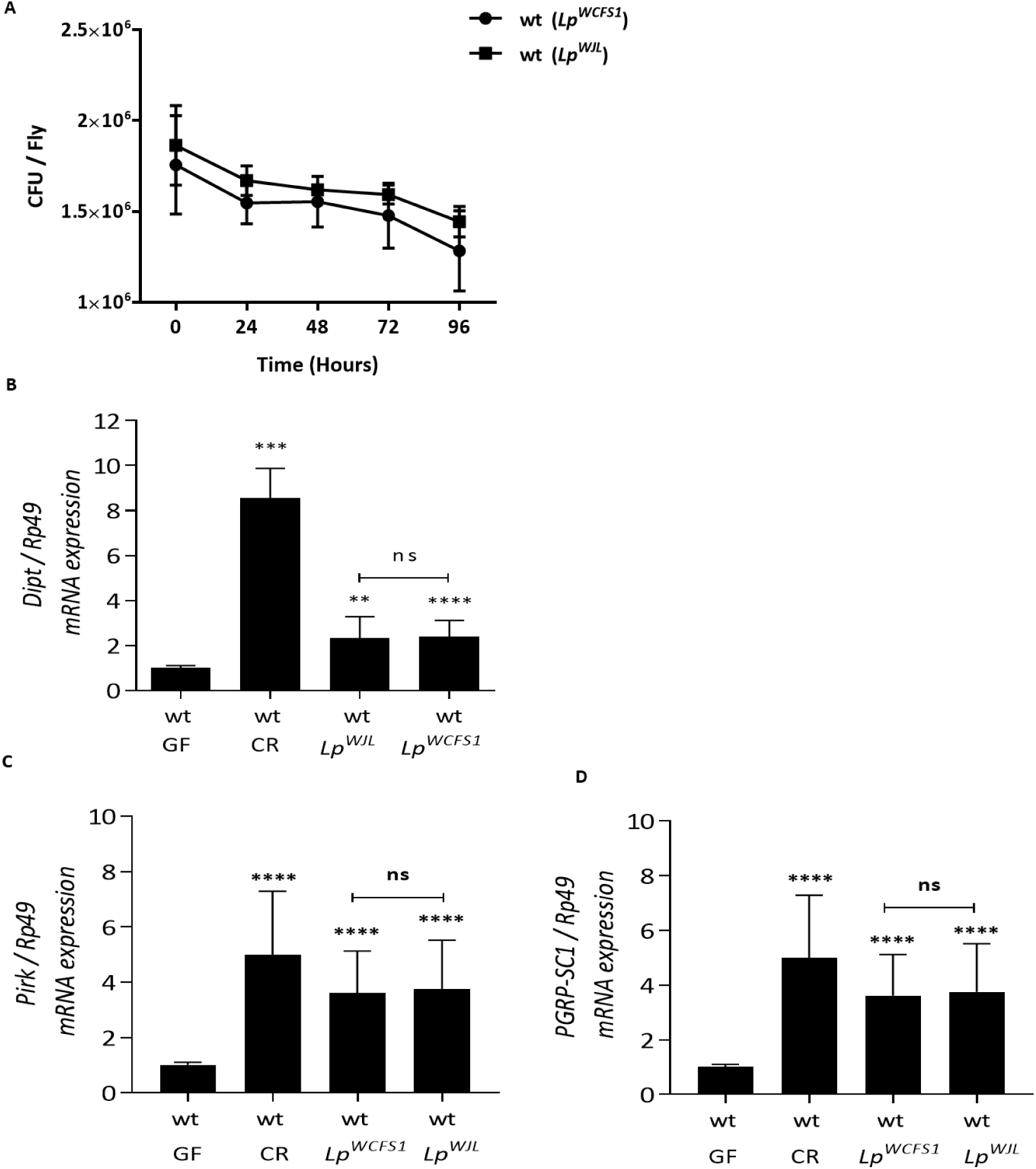
*Lp^WCFS1^* strain shares with *Lp^WJL^* strain the ability to colonize the *Drosophila* gut and to induce the IMD pathway. (A) Internal bacterial loads retrieved from the guts of Germ Free (GF) wild-type (wt) flies fed with *Lp^WCFS1^* or *Lp^WJL^* for three days. Colony Forming Unit (CFU) counting was performed every 24 hours after transferring the monoassociated flies to a fresh cornmeal medium vial. (B, C and D) Relative expression of the *Diptericin* (*Dipt*) (B) *Pirk* (C) and *PGRP-SC1* (D) transcripts in the midgut of Germ Free (GF) wild-type (wt) flies fed with *Lp^WCFS1^* or *Lp^WJL^* for 3 days. Transcripts expression was measured by RT-qPCR in total RNA extracts. Ribosomal protein 49 (Rp49) transcript was used as reference gene. Transcripts levels detected in GF flies is set as a control. Data obtained from three independent experiments are combined in single value (mean ± sd). Statistical tests were performed using the Mann-Whitney test within Prism software (ns: *<u>p</u>* > 0.05; *: 0.01< *p* < 0.05; **: 0.001< *p* < 0.01; ***: 0.0001< *p* < 0.001; ****: *p* < 0.0001).

**Figure 3 – figure supplement 2:**
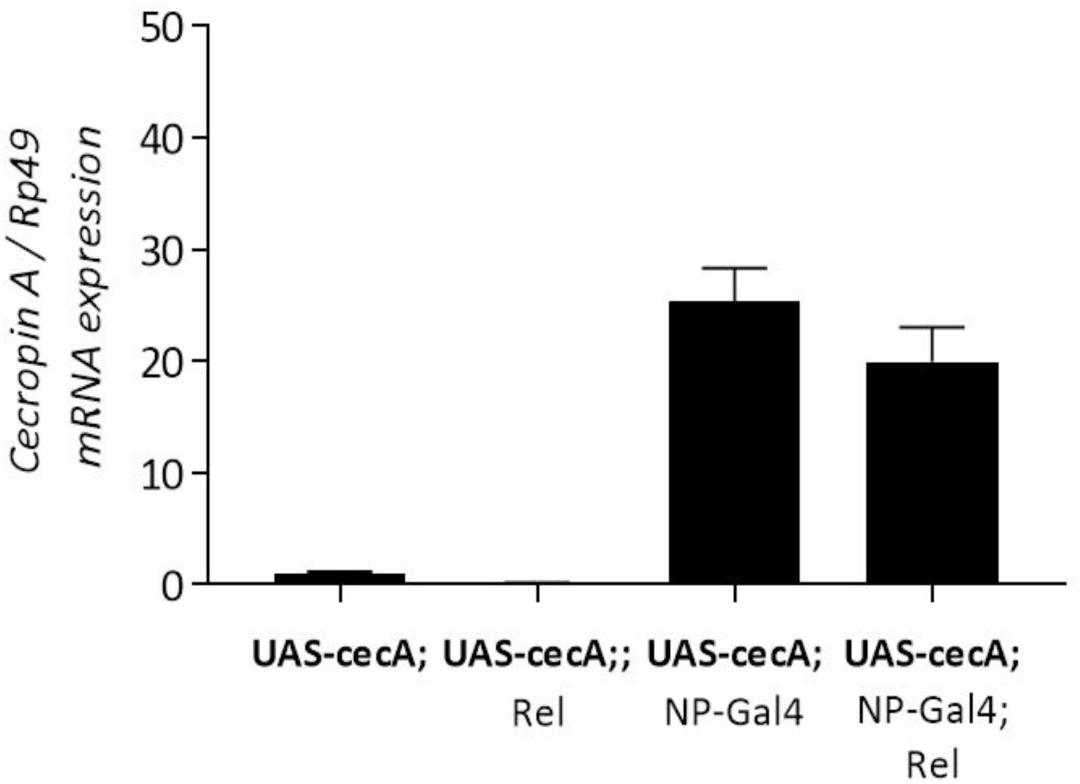
Overexpression of *Cecropin* A gene in the gut of adult flies. The expression of a *cecropin A* transgene (UAS-cec) was driven using a midgut specific driver, NP-Gal4, in wild-type (wt) or *Rel* mutant flies. The *cecropin A* expression a was quantified by RT-qPCR from the RNA extracts of dissected midguts of adult flies following their incubation for 3 days at 29 °C. Wt flies carrying the UAS-cec transgene were used as control. Data obtained from three independent experiments are combined in single value (mean ± sd).

**Figure 3 – figure supplement 3:**
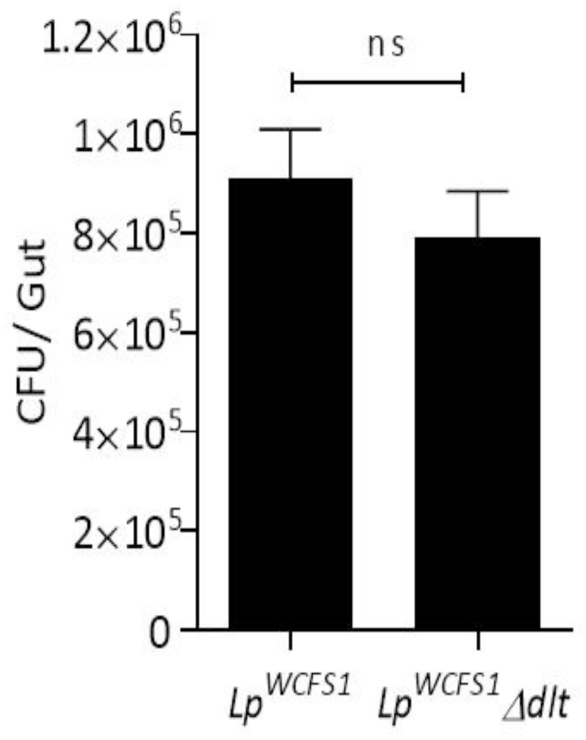
Internal bacterial loads retrieved from the guts of Germ Free (GF) wild-type flies fed with *Lp^WCFS1^* or *Lp ^WCFS1Δdlt^* for three days. Data obtained from three independent experiments are combined in single value (mean ± sd). Statistical tests were performed using the Mann-Whitney test within Prism software (ns: *p* > 0.05)

**Figure 3 – figure supplement 4:**
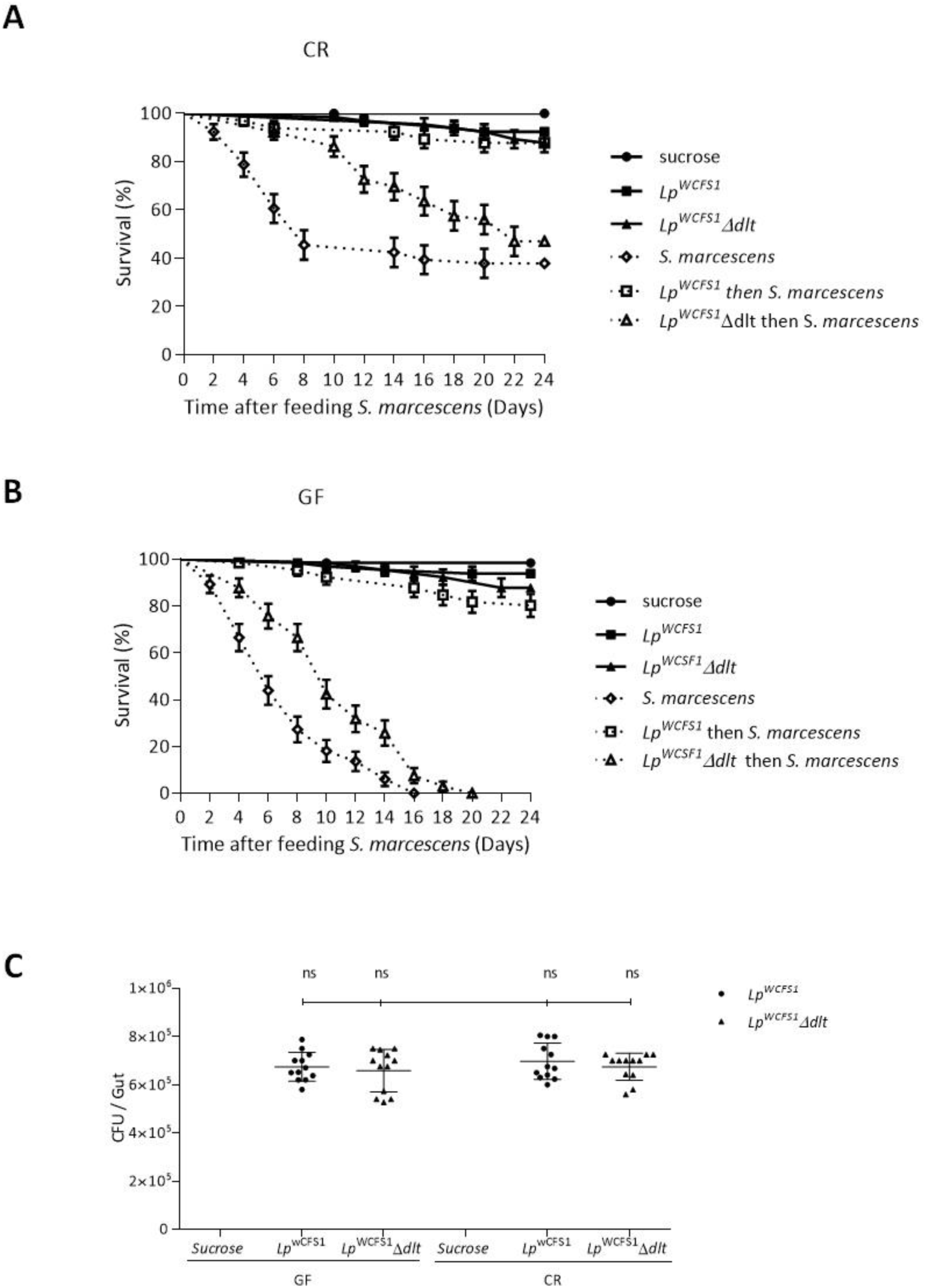
D-alanylation of WTAs are essential for the probiotic effect of *Lp ^WCFS1^* in *Drosophila*. (A and B) Conventionally reared (CR) (A) and Germ Free (GF) (B) wt flies were fed with *Lp^WCSF1^* or *Lp ^WCFS1^Δdlt* for 3 days prior to *S. marcescens* challenge. Flies were fed a sucrose suspension of *S. marcescens* for 2 days and then transferred to a fresh cornmeal vial. (B) Internal bacterial loads retrieved from the guts of Germ Free (GF) or conventionally reared (CR) wt flies fed with *Lp^WCFS1^* or *Lp ^WCFS1^Δdlt* for three days. Data obtained from three independent experiments are combined in single value (mean ± sd). Statistical tests were performed using the Mann-Whitney test within Prism software (ns: *p* > 0.05).

**Figure 5 – figure supplement 1.**
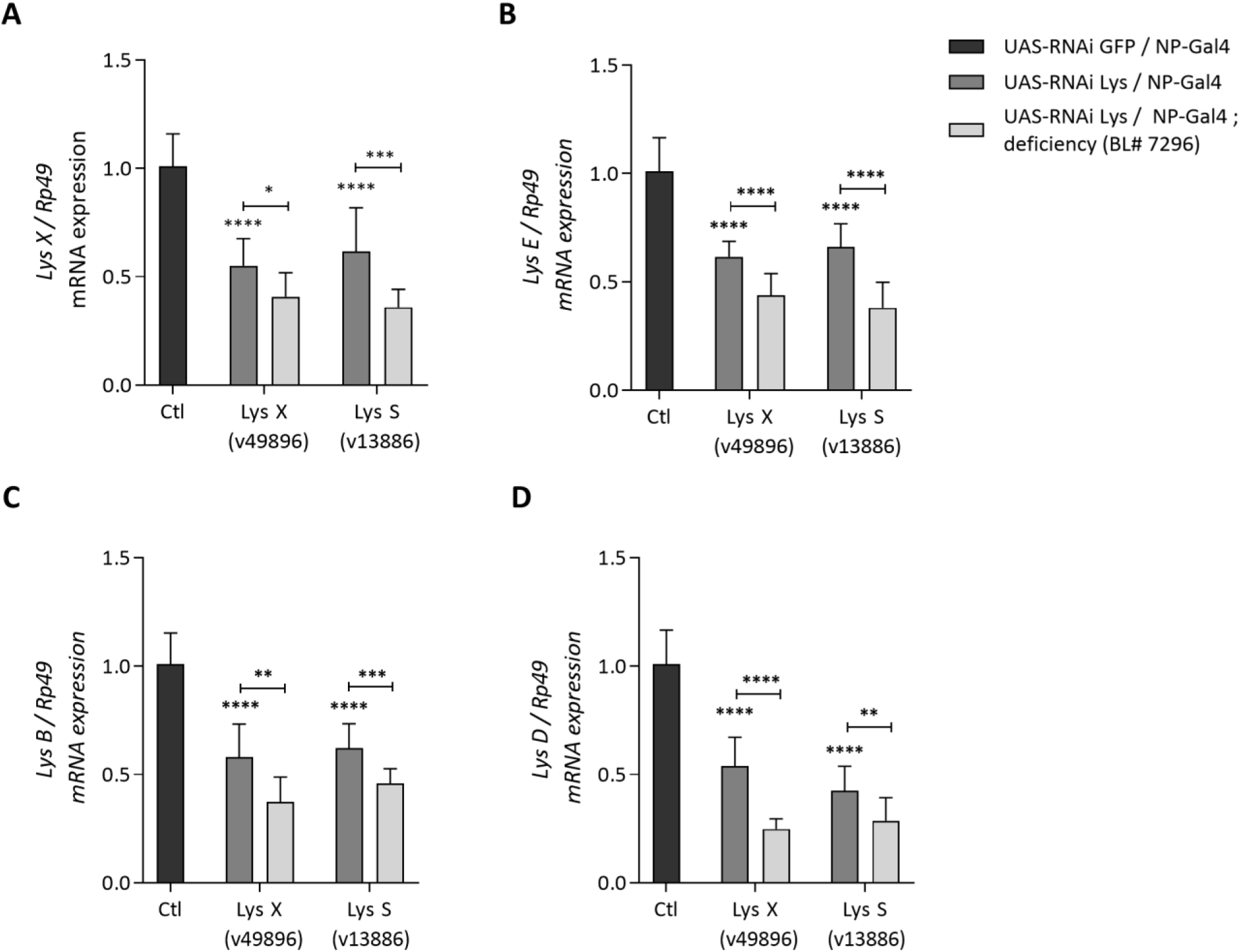
RNAi silencing of *lysozyme* genes in the gut. Silencing of the lysozyme genes was performed by the expression of RNAi constructs (UAS-RNAi) in the gut of adult flies using the midgut specific driver NP-Gal4. Two different transgenic fly lines (v49896) and (v13886) were used. These harbor different RNAi construct targeting the *LysX* or the *LysS* transcripts respectively. Silencing of lysozyme genes is further aggravated by crossing these transgenic flies to a fly line carrying a chromosomal deletion (deficiency BL#7296) uncovering the Lysozyme locus in the *Drosophila* genome. A transgenic line expressing an RNAi construct targeting the GFP transcript is used as a control. Relative expression of the *Lysozymes X* (A), *E* (B), *B*(C) and *D* (D) transcripts is measured in total RNA extracts from the midgut of the transgenic flies of the different genetic background Ribosomal protein 49 (Rp49) transcript was used as reference gene. Transcripts levels are compared to that detected in UAS-RNAi GFP transgenic flies that is arbitrary set to 1. Data obtained from three independent experiments are combined in single value (mean ± sd). Statistical tests were performed using the Mann-Whitney test within Prism software (ns: *p* > 0.05; *: 0.01< *p* < 0.05; **: 0.001< *p* < 0.01; ***: 0.0001< *p* < 0.001; ****: *p* < 0.0001).

## Sources Data

**Figure 1 – source Data 1.** p values for figure 1.

**Figure 2 – source Data 1.** p values for figure 2.

**Figure 3 – source Data 1.** p values for figure 3.

**Figure 4 – source Data 1.** p values for figure 4.

**Figure 5 – source Data 1.** p values for figure 5.

## Additional files

### Supplementary files

Supplementary file 1. List of primers used for the RT-qPCR reaction.

